# Multivariate directed connectivity analysis (MDCA) enables decoding of directed interactions in neural and social networks

**DOI:** 10.1101/263665

**Authors:** Alexander Schlegel, Bennet Vance, Prescott Alexander, Peter U. Tse

## Abstract

Many scientific fields currently face the daunting task of studying the dynamics of complex networks. For example, while we know that the rich mental phenomena of humans and other animals are mediated by complex systems of neural circuits in the brain, the mechanistic links between these biological networks and the functions that they mediate are poorly understood. Here we present a novel class of methods, termed *multivariate directed connectivity analysis*, to investigate network dynamics via patterns of directed interactions between network nodes. We validate these methods using simulated data and apply them to three real-world datasets, two neuroscientific and one investigating the 2016 US presidential candidates’ influence on the social media service Twitter. We find that these methods enable novel understanding of how information processing is distributed across networks. The methods are generally applicable to the study of dynamic information networks in biological, computational, and other fields of research.

## Introduction

Myriad phenomena currently at the forefront of scientific investigation emerge from interactions in complex networks. Examples of such networks can be found in molecular signaling and metabolic pathways in cells (Guimera and Amaral, 2005), the nervous systems of organisms as simple as the nematode *C. elegans* or as complex as humans and other mammals (Sporns, 2014; Towlson et al., 2013), social networks on the Internet (Ahn et al., 2007; Borge-Holthoefer et al., 2015), ecological systems (Montoya et al., 2006; Sugihara et al., 2012), and planetary climate (Mosedale et al., 2006). In particular, the hundreds of trillions of connections among tens of billions of neurons in the human neocortex constitute what may be the most complex network in the known universe (Bassett and Gazzaniga, 2011; Pakkenberg et al., 2003). While neuroscientists are confident that mental phenomena are realized via interactions among such neural connections, empirical investigation of mechanistic links between these interactions and the mind has proven difficult.

Over the last 25 years, new research tools such as functional magnetic resonance imaging (fMRI) (Haxby et al., 2014), multi-electrode recording arrays (Mante et al., 2013), and two-photon imaging (Peron et al., 2015) have allowed brain activity to be measured at unprecedented levels of detail and complexity. These tools have helped to revolutionize our conception of information processing in the brain: While neurons represent a low level unit of information processing, higher level representations and processes may emerge fundamentally at multiple levels and timescales of *interaction* between hierarchical networks of these units. Population coding (Mante et al., 2013), distributed representational spaces (Haxby et al., 2014), and cortex-wide functional networks (Schlegel et al., 2016; Turk-Browne, 2013) are examples of this interaction-based conception of information.

Recent promising work has combined these new measurement tools with graph theoretical measures to describe neural interactions at a “connectome”-wide level of analysis. But this bird’s eye view of the brain leaves our understanding of the functional roles of specific network connections murky at best (Bullmore and Sporns, 2009; Hagmann et al., 2008; Zuo et al., 2012). Measures have also been developed to analyze multivariate information carried by brain-wide functional connectivity patterns (Richiardi et al., 2011; Shirer et al., 2012), but these methods do not resolve the information processing roles of specific network nodes, nor the direction of their influence. A full understanding of any information processing network requires a description of the mechanistic roles played by specific, directed network interactions.

Several analytical measures have been developed to investigate directed influence between the nodes of information networks. Some measures, such as Granger-causality (Barnett and Seth, 2014) and transfer entropy (Lizier et al., 2011), define influence via the statistical ability of the past of one signal to predict the future of another signal. Other measures such as dynamic causal modeling (Friston, 2011) and convergent cross mapping (Sugihara et al., 2012) attempt to move beyond statistical prediction by uncovering the underlying causal structures in a network. However, each of these measures exhibits two properties that limit its usefulness in understanding the functions of specific interactions in a complex network:

First, existing applications of these directed connectivity (DC) measures are concerned primarily with determining whether one node exerts influence over another. This is an important question to ask of networks that exhibit relatively sparse, well-organized connection profiles. However, an emerging view of the brain as a densely connected, highly distributed network makes this approach less relevant, since the nodes in such a network will likely exert a complex pattern of control over many if not all other nodes. More relevant to understanding the behavior of complex networks such as these would be to study the specific information processing contributions of network node pairs.

Second, much like the univariate analyses that dominated the early fMRI literature, existing DC measures can detect coarse changes in the univariate level of influence between nodes but are insensitive to subtle, complex patterns of connectivity that may differ between the states of highly dynamic networks (Haxby et al., 2001). Even nominally “multivariate” DC measures such as multivariate Granger-causality (MVGC) (Barnett and Seth, 2014) and multivariate transfer entropy (MVTE) (Lizier et al., 2011) still yield only a single scalar quantity to characterize the directed influence between two (multivariate) spaces. Important questions can be asked using these existing methods; for example, whether cortico-cerebellar connectivity is modulated by the difficulty of a visuomotor task (Lizier et al., 2011). However, such methods would in general be unable to resolve information-based questions; for example, if multivariate *patterns* of cortico-cerebellar connectivity can be used to decode whether a patient is typing or playing the piano. Answering the latter kind of question may prove more important for the development of technologies such as brain-computer interfaces (Nicolelis, 2001).

To address the need for information-based approaches to network interactions, here we develop and validate a new class of methods, termed *multivariate directed connectivity analysis* (MDCA), that extend existing DC measures by allowing informational differences to be decoded in specific directed network interactions. Figure 1 presents a schematic overview of MDCA. First, two multidimensional spaces (or “nodes”) within a network—labeled the *source* and the *destination*—are defined, the goal being to evaluate information processing carried out via directed connectivity from the source to the destination. Examples of such spaces are regions of interest (ROIs) in fMRI data, electrode groups in electroencephalogram (EEG) or electrocorticogram (ECoG) data, or groups of users in a social media network. Because these two spaces are multidimensional, the interactions between them have a multivariate structure. Thus, we may use any existing DC measure (e.g. Granger-causality or transfer entropy) to compute the directed influence from every component of the source space to every component of the destination space. This process yields a pattern (or “graph”) of connectivity between the source and destination, and multiple instances of these patterns can be computed and labeled (e.g. one for each condition and run of an fMRI experiment). This treatment of source-destination interactions as multivariate then creates the potential for a large set of multivariate methods to be applied in order to probe the information associated with those interactions. For example, multivariate classification between experimental conditions can be performed using source-destination connectivity as the input pattern (Norman et al., 2006), or representational similarity analysis can evaluate the relationship between source-destination connection profiles over a range of behaviors (Kriegeskorte et al., 2008). A key advance of this approach is that it enables the measurement of differences in the pattern of directed connectivity between two spaces, *even when the overall direction and magnitude of connectivity remain fixed*. As an analogy, one might consider telecommunications cables that transmit roughly the same amount of data at any instant, while the informational content of those data transmissions changes considerably from moment to moment.

**Fig. 1.**
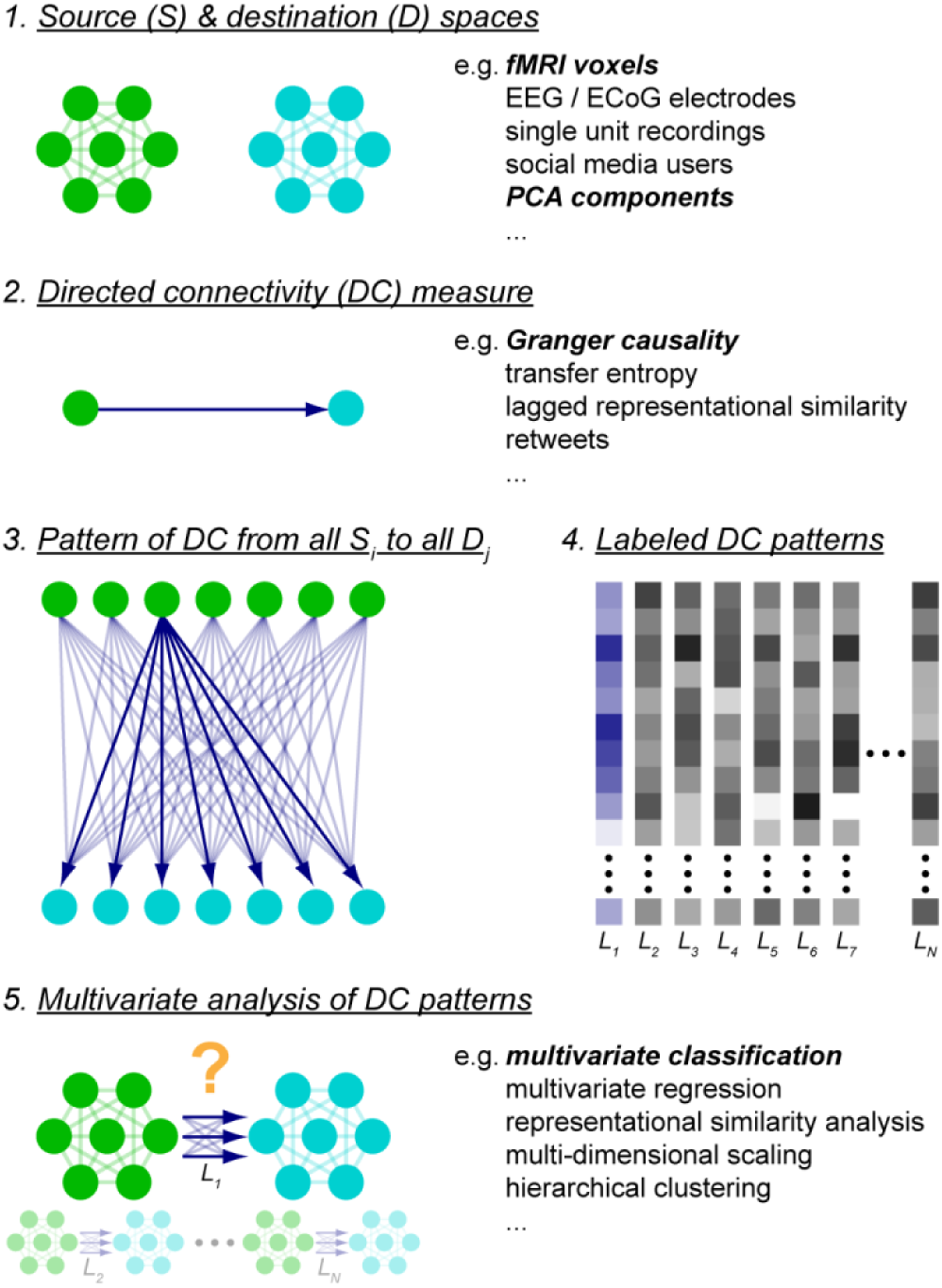
General schematic of multivariate directed connectivity analyses. *1.* MDCA begins with the definition of two multidimensional spaces, a source and a destination (green is the source and cyan is the destination). *2.* Any of several existing directed connectivity measures is chosen to compute the directed influence of each process in the source space (S_i_) on each process in the destination space (D_j_). *3.* This computation yields a multidimensional pattern of directed connectivity (with one dimension for each pair of source and destination processes, S_i_ and D_j_). *4.* A set of these computed patterns are labeled (e.g. L_k_) according to experimental conditions. Each column represents a single labeled pattern. *5.* This procedure then enables any number of existing multivariate analyses (e.g. classification, representational similarity analysis) to be performed on these directed connectivity patterns.

Below, we first simulate data for a standard fMRI experiment to validate MDCA as a method that can resolve differences between experimental conditions to which several other standard analytical methods are insensitive. We then apply MDCA to three real-world datasets—one from an fMRI study of the manipulation of visual imagery, one from an EEG study of action preparation, and a final dataset measuring the influence of the 2016 Republican U.S. presidential candidates on the social media service Twitter. The results of these three analyses demonstrate the wide ranging potential for application of MDCA. Our fMRI analysis provides evidence that highly distributed processing mediates the manipulation of visual imagery, arguing against standard, anatomically modular models of working memory (Baddeley, 2003; Postle, 2006). Our EEG analysis suggests that distributed patterns of processing between posterior and anterior cortical regions contain rich information about the preparation and execution of actions and may therefore be useful in the development of brain-computer interfaces (Nicolelis, 2001). Finally, our Twitter dataset demonstrates that MDCA’s potential applications extend beyond the brain into a range of other complex networks such as social interactions on the Internet.

## Materials & Methods

### Simulated fMRI Dataset

#### Data generation

Here we simulated an experiment in which 15 subjects each completed 10 runs of a block-design fMRI experiment. In each run, 10 TR blocks of two experimental conditions were presented in counterbalanced order, interleaved with 5 TR rest periods, with five repetitions of each condition during the run.

The simulated data for this experiment consisted of fMRI time courses collected with 2-second TR measured from two 100-voxel regions of interest (ROIs). Crucially, these data simulate a situation in which information processing in two brain regions differs between the two experimental conditions, but in a way that existing methods would not detect. Specifically, mean activity and multivariate activity patterns in each region and even mean directed connectivity between the two ROIs do not differ significantly between the two conditions. Thus, existing univariate, multivariate, and directed connectivity measures would find no differences in the ROIs between conditions. However, *patterns* of directed connectivity between the two regions do differ systematically between the two conditions. This type of behavior could be expected in networks that continuously mediate an array of complex processes differing only in the kind of information processing rather than the amount of information processing that occurs within and between nodes. Our question here is: Can MDCA resolve these pattern-level differences and thus find evidence that the two ROIs interact to process the two conditions differently?

Data were first generated in a hypothetical functional space before being mixed linearly into the voxel space that would be measured by an fMRI scanner. Each functional space time course was generated according to the following equations:

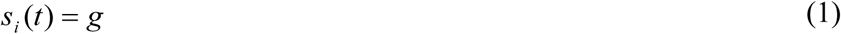

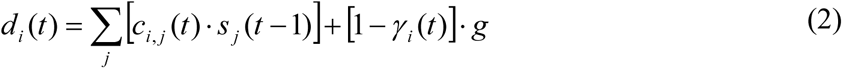

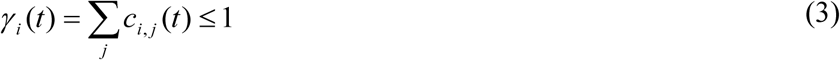

where *s(t)* and *d(t)* are the values of the multidimensional source and destination signals at integer sampling time *t* (in TRs), *s_i_(t)* is the *i*th component of *s(t), g* is a Gaussian noise process, *d_i_(t)* is the *i*th component of *d(t), c_i,j_(t)* is the *i,j*th component of the DC matrix *C(t)* which defines the pattern of directed influence by the source over the destination at time *t*, and *γ_i_(t)*—the sum of the *i*th row of *C(t)*—controls the relative influence of the source signals and the Gaussian process to the variance in the *i*th destination signal (i.e. higher values of *γ* lead to a greater causal coupling strength between the source and destination). Note that the value of *c_i,j_(t)* determines the influence of the *j*th source signal at time *(t – 1)* on the *i*th destination signal at time *t*.

In this simulation, *C(t)* alternates between three values according to the block design in Figure 2A: *C_A_* (active during blocks of condition A), *C_B_* (active during blocks of condition B) and *C_rest_* (active during rest periods). Each DC matrix is sparse so that only a subset of source processes exerts influence over only a subset of destination processes, while the remaining processes are not causally linked. Here 10 of the 100 source processes are potentially linked to 10 of the 100 destination processes (i.e. *c_i,j_*(*t*) = 0 for *i* > 10 or *j* > 10), with the additional constraint that ¾ of the remaining values are 0. This constraint was used in order to simulate a situation in which each causally linked source process was linked to some, but not all, of the causally linked destination processes. As a final constraint, the non-zero values of each DC matrix are generated randomly while holding *γ_i_(t)* constant for all values of *i* and *t*. Because of this constraint on the DC matrices, the source space exerts a constant magnitude of influence over the destination space throughout the experiment, while only the pattern of this influence depends on the condition. Note that the only aspect of the data that differs between conditions is the DC matrix *C_K_* that is active during each of those conditions.

**Fig. 2.**
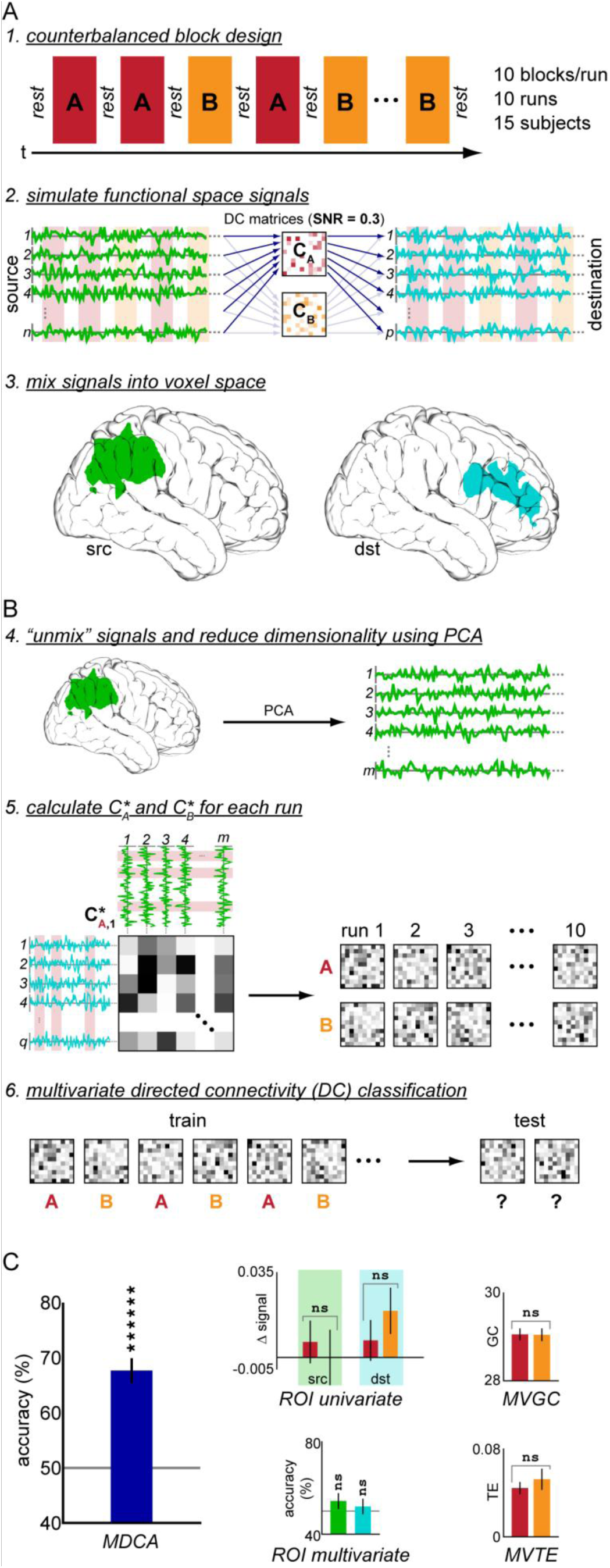
Simulated experiment demonstrating sensitivity of MDCA. **A.** Generation of simulated data. *1.* The experiment uses a standard fMRI block design in which two conditions are presented with interleaved rest periods. *2.* We simulate a neural source (green) and destination space (cyan), each consisting of a set of functional processes (univariate time signals). Processes in the source space influence those in the destination space according to one of two patterns of directed connectivity (C_A_ during blocks of condition A and C_B_ during blocks of condition B). *3.* These functional space data are mixed linearly into two voxel-based regions of interest (ROIs) that represent what would be measured in an fMRI experiment. **B.** Data analysis. *4.* Each measured signal is concatenated across runs and principal components analysis is performed on each ROI independently in order to partially recover the functional space data and to control the dimensionality of the space. *5.* Estimates of the original directed connectivity patterns (C_A_* and C_B_*) are constructed for each run and condition by calculating the Granger-causality from each source signal to each destination signal. *6.* A standard multivariate classification analysis tests for differences in directed connectivity patterns between the two conditions. **C.** Analysis results. The directed connectivity classification (MDCA) detects significant differences between the two conditions, whereas none of four standard methods (ROI univariate analysis, ROI multivariate classification, multivariate Granger-causality, and multivariate transfer entropy) are sensitive to these differences. Error bars are standard errors of the mean across subjects. ******: p ≤ 10^−6^.

The signal-to-noise ratio (SNR) of the data was controlled via the ratio of the total power of the causally-linked signals to that of the non-linked signals. The magnitudes of the non-zero values of *C*(*t*) were scaled to give a causal coupling strength of 0.5, with coupling strength defined as the ratio of the variance in the destination signals contributed by the source processes to the variance contributed by the Gaussian processes. In other words, the random Gaussian processes had twice as much influence over the causally linked destination signals as did the source processes. We additionally scaled the magnitudes of the signals in order to achieve a signal-to-noise ratio (SNR) of 0.3, with SNR defined as the ratio of the total variance of the 10 causally linked signals to the total variance of the 90 unlinked signals.

Finally, the functional space signals generated as described above were mixed linearly, using randomly generated matrices of mixing weights, into voxel-based source and destination ROIs that represent the data that would actually be measured in an fMRI experiment.

#### Data analysis

For each simulated subject, data for each ROI were first concatenated across runs and then analyzed in one of five different ways, as follows:

##### Univariate ROI analysis

For each subject and condition, the mean signal amplitude across voxels and runs was computed, and the difference in these mean amplitudes between conditions was evaluated using a paired t-test across subjects.

##### Multivariate ROI classification

For each subject, run, condition, and ROI, a voxel-based pattern was constructed by computing the mean signal amplitude of each voxel during that condition. For each subject and ROI, we then performed a standard multivariate classification analysis using leave-one-run-out cross-validation in which we tested whether a machine classifier (here a linear support vector machine with C = 1) could be trained to distinguish between conditions A and B based on the voxel-based patterns constructed for each run, condition, and ROI. A group-level one-tailed t-test then compared the subject-wise classification accuracies to chance, which was 50%.

##### Univariate directed connectivity analysis

Data were analyzed as in the multivariate directed connectivity classification (described below), except that instead of computing GC-graphs, either the multivariate Granger-causality (MVGC) (Barnett and Seth, 2014) with lag = 1 TR or multivariate transfer entropy (MVTE) (Lizier et al., 2011) using a Kraskov estimator with K = 4 and lag = 1 TR was calculated from the source ROI to the destination ROI for each subject and condition. These scalar values were then compared between conditions using a paired t-test across subjects.

##### Multivariate directed connectivity classification

(see Figure 2B). For each subject, concatenated data were transformed using principal components analysis, and all but the top 10 components from each ROI were discarded. We performed this transformation for two reasons: First, it attempts to “unmix” the measured voxel-based signals in order to recover a space similar to the original functional space in which each component represents a statistically separable process. Second, it allows us to control the dimensionality of our source and destination spaces and thus the size of the estimated DC patterns we will construct.

For each run and condition, a new signal was then constructed for each ROI by extracting and concatenating data from that run and condition only. A 1 TR time lagged version of this signal was also constructed, and these signals were used to compute a graph of Granger-causality (GC-graph) from each source signal to each destination signal. Granger-causality is a method for measuring directed functional connectivity between signals that has been shown to be a valid and powerful tool for analyzing a range of data modalities, including fMRI (Barnett and Seth, 2014; Brovelli et al., 2004; Cadotte et al., 2008; Keil et al., 2009; Sato et al., 2006; Seth, 2010; Wen et al., 2013). This process yielded a graph of Granger-causality (GC-graph) from the source to the destination for each subject, run, and condition.

A multivariate classification analysis as described above was then performed to test whether these GC-graphs could be used to distinguish between the two experimental conditions.

#### SNR plots

The plots in Figure 3A–D were generated by running multiple simulations while varying the SNR and one other parameter (either number of subjects, number of runs, number of data samples per DC pattern estimation, or causal coupling strength between source and destination). For each tested SNR value, each plot shows the best-fit estimate of the parameter value needed to achieve a significant DC classification result (p ≤ 0.05).

**Fig. 3.**
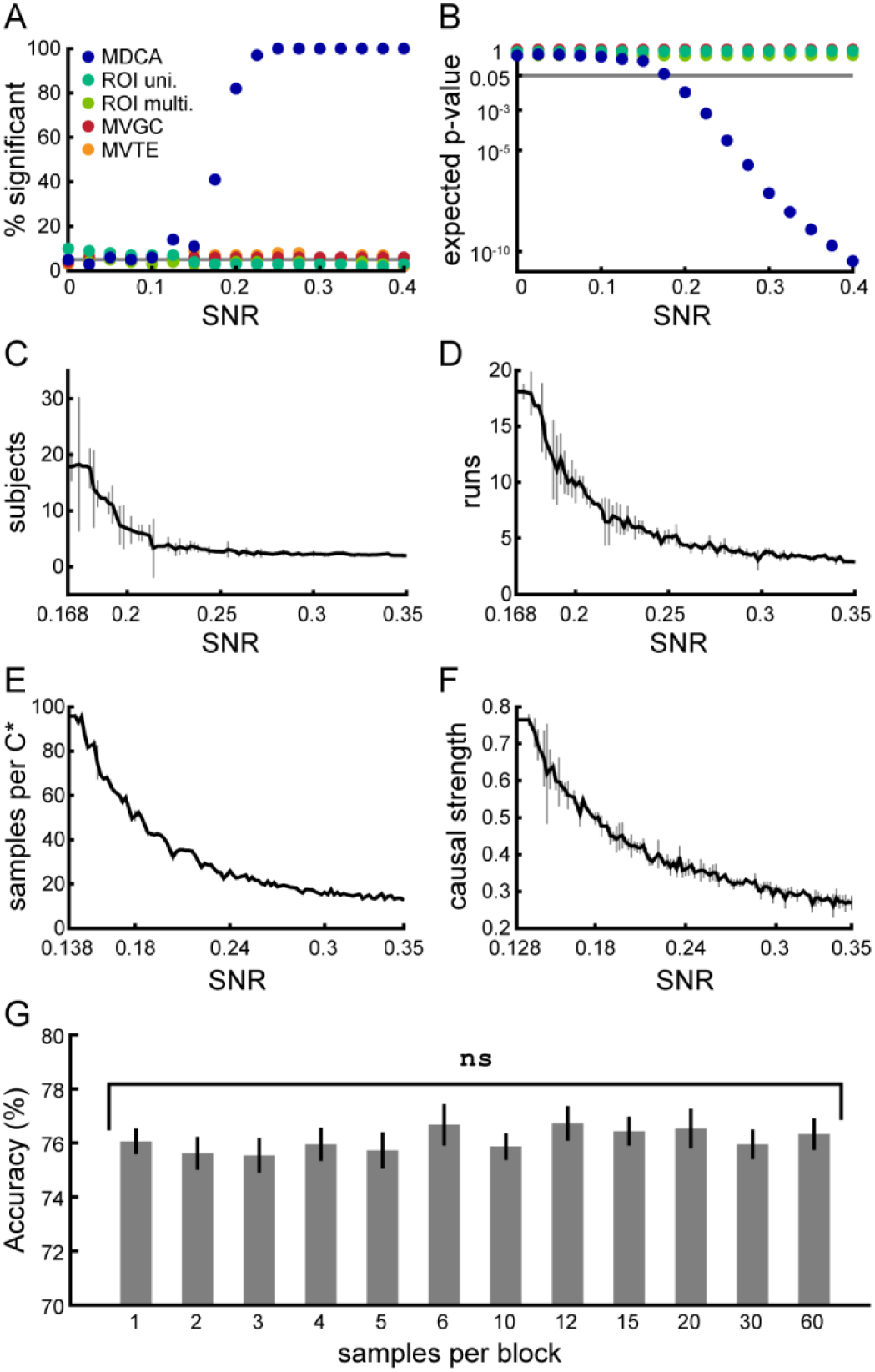
Sensitivity of MDCA. Unless specified, all parameters are the same as those used in the stimulation from Figure 2. **A.** The simulation from Figure 2 was run 100 times at each of a range of SNR values (see text for definition of SNR). This plot shows the percentage of simulations that led to significant results for each of the five analyses. Only MDCA rises above the expected false positive rate of 5%, starting at around SNR = 0.15. **B.** For the same simulations from panel A, the expected *p*-value of each analysis. Expected *p*-value for a given analysis and SNR is the *p*-value corresponding to the mean effect size of that analysis across all simulations at that SNR. MDCA reliably reaches significance starting at around SNR = 0.2, while no other analysis reaches threshold for any SNR. **C.** The simulation from Figure 2 was run repeatedly at a range of SNR values to determine the minimum number of subjects needed in order to achieve significant results (expected p ≤ 0.05). Error bars are 50% confidence intervals over the simulations run for each SNR value. **D.** Same as C, but for number of runs. **E.** Same as C, but for number of samples per GC-graph estimate. **F.** Same as C, but for causal strength (see text). **G.** Results of analysis comparing classification accuracy for a range of block durations, with number of samples per GC-graph estimate held constant. Error bars are standard errors of the mean across 30 simulations per block duration value.

### Real fMRI dataset

#### Participants

40 participants (29 females, aged 18–32 years) with normal or corrected-to-normal vision gave informed written consent according to the guidelines of the Committee for the Protection of Human Subjects (CPHS) at Dartmouth College prior to participating (IRB #15822). Participation consisted of two 1.75 hour fMRI scanning sessions.

#### Experimental design

We replicated the protocol used in Schlegel et al. (2016). Participants completed 15 fMRI runs, each of which consisted of 16 trials interleaved with 8 sec. of rest. In each trial, participants performed one of four mental operations on one of four abstract visual shapes. The four mental operations were: 90°Clockwise rotation, 90°Counterclockwise rotation, horizontal flip, and vertical flip. The four abstract shapes are shown in Figure 4A: two shapes were constructed from a 4×4 rectangular grid, and two were constructed from an analogous polar grid. To equate the visual presentation between conditions, we did not display the shape or operation to use in a given trial. Instead, each shape was mapped to one of the letters A, B, C, or D, and each operation was mapped to one of the numbers 1, 2, 3, or 4. At the start of each trial, a 2-second-long prompt screen displayed four letter/number pairs (e.g. “C3”). An arrow pointed to one of these pairs to indicate the shape and operation to use for the current trial. This screen was replaced by a fixation dot for 6 sec. during which the participant performed the indicated mental operation on the indicated shape. After this period, a 2-second-long test screen displayed each of the four shapes at various orientations relative to the starting orientations learned by the participants. The participant was instructed to identify the current trial’s shape on the screen and indicate via a button press within that 2 sec. period whether it was in the orientation that would result from the trial’s indicated operation. In half of the trials the shape was in the correct orientation, and in the other half it was in a random, incorrect orientation. Operations and shapes were counterbalanced across all trials, and correct/incorrect trials and display positions were randomized. In order to encourage attentiveness, participants were paid based on their performance (receiving money for correct responses and losing money for incorrect responses, with a minimum base rate of reimbursement). Each stimulus and operation occurred four times per run (60 times in total during the experiment), and 240 trials were administered over each scanning session.

**Fig. 4.**
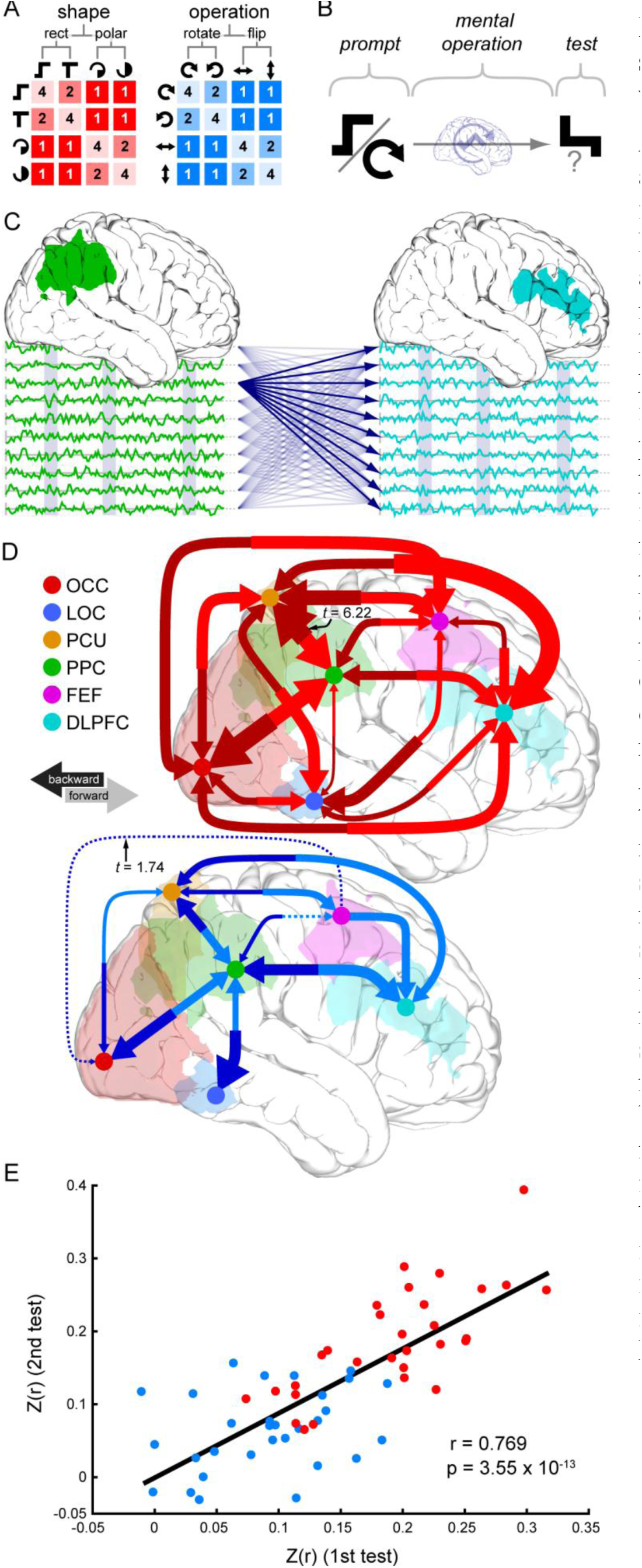
Data from a real-world fMRI study of the mental manipulation of visual imagery. **A.** Participants imagined one of four abstract shapes and performed one of four mental operations on that shape. Both shapes and operations are related in a two-level hierarchy of similarity, represented in matrix form below each hierarchy. **B.** On each trial, the participant was prompted with a shape to imagine and a mental operation to perform, was then given six seconds to perform the operation on the shape, and finally indicated whether a test stimulus matched the correct output of the prompted operation. **C.** Data for the analysis here were DC patterns between pairs of cortical ROIs. **D.** Results of DC classification analysis between each directed pair of ROIs. Each arrow represents a successful classification. Arrow width represents effect size (see text). All results are FDR corrected for multiple comparisons. Red arrows are successful classifications of mental representation, and blue arrows are successful classifications of mental manipulation. **E.** Correlation of analysis results between the first and second sessions. Red dots are from the mental representation classification analysis, and blue dots are from the mental manipulation classification analysis. *Z*(*r*) is the Fisher’s *Z*-transformed correlation between the model similarity structure from Figure 4A and the confusion matrix from the four-way classification analysis for a given directed pair of ROIs. MDCA shows a high test-retest reliability [*r* = 0.769, *t*(58) = 9.16, *p* = 3.55×10^-13^].

#### MRI acquisition

MRI data were collected using a 3.0-Tesla Philips Achieva Intera scanner with a 32-channel sense head coil located at the Dartmouth Brain Imaging Center. One T1-weighted structural image was collected using a magnetization-prepared rapid acquisition gradient echo sequence (8.176ms TR; 3.72ms TE; 8°Flip angle; 240×220mm FOV; 188 sagittal slices; 0.9375×0.9375×1mm voxel size; 3.12 min acquisition time). T2*-weighted gradient echo planar imaging scans were used to acquire functional images covering the whole brain (2000ms TR, 20ms TE; 90°Flip angle, 240×240mm FOV; 3×3×3.5mm voxel size; 0mm slice gap; 35 slices).

#### MRI data preprocessing

High-resolution anatomical images were processed using the FreeSurfer image analysis suite (Dale et al., 1999). Standard preprocessing of fMRI data was carried out: data were motion and slice-time corrected, high pass filtered temporally with a 100-s cutoff, and smoothed spatially with a 6mm full-width-at-half-maximum Gaussian kernel, all using FSL (Smith et al., 2004). Data from each run were concatenated temporally for each participant after aligning each run using FSL’s FLIRT tool and demeaning each voxel’s time course. Data were then prewhitened using FSL’s MELODIC tool (i.e. principal components were extracted using MELODIC’s default dimensionality estimation method with a minimum of 10 components per ROI).

#### fMRI analysis

The six regions of interest (ROIs) were defined as in Schlegel et al. (2016) (PPC [posterior parietal cortex], PCU [precuneus], LOC [lateral occipital cortex], DLPFC [dorsolateral prefrontal cortex], and FEF [frontal eye fields] were defined functionally, while OCC [occipital cortex] was defined anatomically). For each directed pair of ROIs, GC-graphs were estimated as in the simulation, using the first 5 TRs of each correct-response trial, shifted by 1 TR to account for the hemodynamic response function (HRF) delay. Note that, while HRFs typically take longer to reach their peak, shifting by only 1 TR here allowed us to include more data that could potentially show condition-specific effects of interaction between areas. The exact shift used is not crucial, since including too much data would only decrease our effective SNR, and multivariate classification methods are in general effective at separating out signal from noise. The most important consideration here is to make sure that the period of signal used to construct the GC-graphs could not be contaminated by data from other conditions.

We used the multivariate classification analysis described above for the simulation. We performed the analysis twice for every directed pair of nodes in the network: once with trials labeled based on shape, and once with trials labeled based on operation. Our input data to the analysis were the GC-graphs estimated separately for each condition and run using the PCA-transformed data of a given pair of ROIs and a lag of 1 TR (2 sec). Figure 4C shows a schematic of one GC-graph for the PPC to DLPFC connection. Because of the hierarchical relationships among our shapes and operations, we evaluated the classifier using a variant of a representational similarity analysis in which our measure of classifier performance was the Fisher’s Z-transformed correlation between the confusion matrices derived from the cross-validation procedure and the model similarity structures from Figure 4A. This measure is more sensitive than classification accuracy because it allows one to take advantage of both correct and incorrect predictions by the classifier and therefore to test for similarities in informational structure between the neural data and the experimental task itself. A higher Z-value indicates a greater match between the task structure and the informational structure of the communication between the two tested regions. For each participant, shape or operation labeling scheme, and directed pair of ROIs, this cross-validation procedure yielded a single Fisher’s Z-transformed correlation. A one-tailed t-test then compared these Z values to 0 across participants. See Schlegel et al. (2016) for more details of this approach to multivariate classification.

### EEG dataset

#### Participants

9 participants (3 females, aged 19–28 years) with normal or corrected-to-normal vision gave informed written consent according to the guidelines of the Committee for the Protection of Human Subjects (CPHS) at Dartmouth College prior to participating (IRB #22950). Participation consisted of a 1.5 hour EEG session.

#### Experimental design

The experiment used a variation of a study by Donchin and colleagues to investigate action preparation (Donchin et al., 1972). Each participant completed four blocks of trials. In each trial, a rapid serial visual presentation (RSVP) stream of letters appeared. The letters were initially blue in color and gradually filled in to become yellow over the 3 second “warning” period. At the end of this period, an imperative stimulus was presented that instructed the participant to perform an action. In block one, the “Go” block, participants knew that the imperative stimulus would always instruct them to move. Participants were paid based on their reaction time. In block two, the “Go/No-go” block, participants were instructed either to move (“Go”) or to withhold a movement (“No Go”). They were paid based on their reaction time and lost half of their earnings above a baseline amount if they responded before the imperative stimulus or during a no-go trial. In block three, participants merely anticipated a stimulus without an associated motor task. In this “Predict” block, participants predicted before each trial began whether the imperative stimulus would be the letter “L” or the letter “R”. They indicated their prediction with a button press and then merely watched the RSVP stream.

Participants were paid only if they guessed correctly. In the fourth “Compute” block, the imperative stimulus was a green or red number that participants were instructed to add to or subtract from, respectively, a running total that they maintained in working memory. Participants were paid based on the difference between their reported total and the true total. Each block consisted of 80 trials, totaling 320 trials during the session.

#### EEG acquisition and preprocessing

EEG data were recorded at 2048 Hz using BioSemi Ag/AgCl high impedance active electrodes located at 32 scalp locations according to the 10–20 system (Jasper, 1958) and on the left and right mastoids. Scalp data were referenced offline to the average mastoid signal, bandpass filtered from 0.1–150 Hz, discrete Fourier transform filtered to remove line noise at 60, 120, and 180 Hz, and time locked to the imperative stimulus. Each epoch was then demeaned, and epochs in which the signal amplitude of either channel FP1 or FP2 exceeded 8μV were discarded.

#### EEG analysis

*Multivariate directed connectivity classification.* For a given window start time and time lag (see Figure 5C), DC patterns were estimated for each trial using Granger-causality and a 250ms window of data. The two spaces for DC pattern estimation were a group of the 10 most posterior EEG electrodes (Oz, Pz, O1, PO3, P3, P7, O2, PO4, P4, and P8) and a group of the 10 most anterior EEG electrodes (AF3, F3, F7, FC1, FC5, AF4, F4, F8, FC2, and FC6). Each group served as both the source and the destination in separate multivariate classification analyses between the four trial types, carried out as described for the simulation above.

**Fig. 5.**
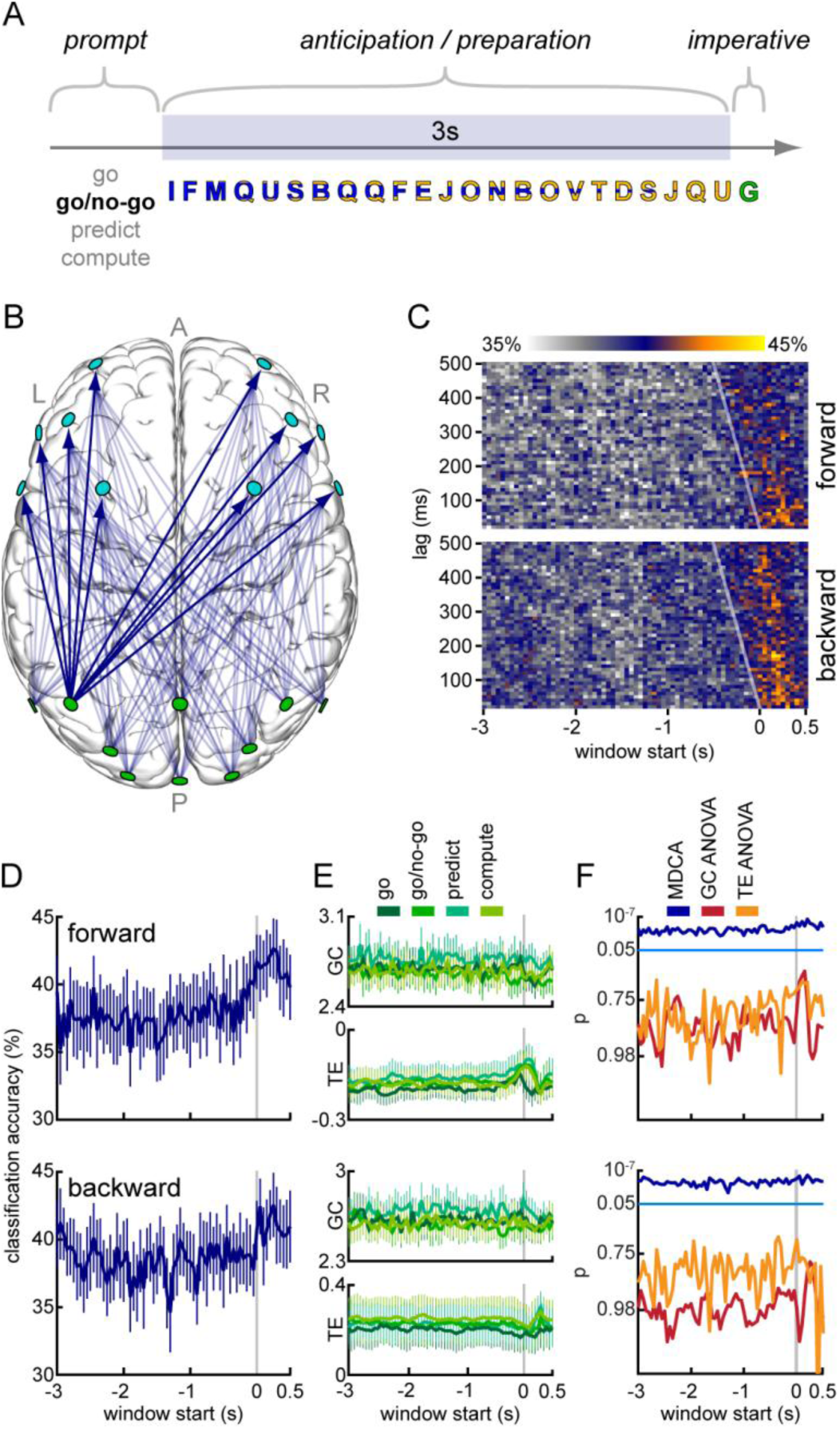
Results from a real-world EEG study of action preparation. **A.** Experimental design. On each trial, an RSVP stream of blue letters gradually filled in to yellow over three seconds. During this time the participant prepared to perform one of four actions. The participant was then prompted by an imperative stimulus to perform the action. **B.** Data for the analysis here were DC patterns between a group of the 10 most posterior and a group of the 10 most anterior EEG electrodes. **C.** Results of DC classification analyses for both forward and backward connectivity. 2D plots show classification accuracy as a function of window start time and lag value used in the Granger-causality calculation. t = 0 at imperative stimulus. Diagonal white line in each plot separates analyses that include data at or after the imperative from those that do not. **D.** Time courses of classification accuracy for lag fixed at 50ms and for forward and backward connectivity analyses. Chance is 25%. Error bars are standard error of the mean across participants. **E.** Time courses of multivariate Granger-causality (MVGC) and multivariate transfer entropy (MVTE) for each connectivity direction and experimental condition. **F.** Log-transformed plots of time courses of p-values for each of the three analyses (DC classification [dark blue], one way ANOVA using MVGC [red], and one way ANOVA using MVTE [orange]). Light blue line is at the significance threshold (p ≤ 0.05).

In an initial analysis we explored how our classifier performed over a range of 250ms time windows starting at the beginning of the preparation period of the trial and continuing in 50ms increments until 500ms after the imperative stimulus had appeared. We also performed the analysis using a range of lag values for the Granger-causality calculation, in order to test whether the influence of the source space over the destination space peaked at particular time lags.

#### Univariate directed connectivity analyses

For a fixed lag of 50ms, either MVGC or MVTE was calculated for each window and trial. These values were then averaged across each condition and participant, and one-way ANOVAs evaluated whether the univariate directed connectivity differed significantly between conditions at each time point. Analyses were performed using both posterior-anterior connectivity and anterior-posterior connectivity.

### Twitter dataset

Twitter is an Internet-based social media service that allows users to follow other users in order to engage in dialog on a massive scale. Not only has Twitter become highly influential in shaping public dialog, it also provides a rich measure of social interaction on a national and worldwide scale (Borge-Holthoefer et al., 2015). In particular, the service has begun to play a significant role in U.S. presidential campaigns. The dataset used here considered the influence of 15 2016 Republican U.S. presidential candidates on the social media service Twitter. User names of each of the candidates are shown in Table S7.

The Twitter API (https://dev.twitter.com/overview/api) was used to collect information about the activity of the candidates and their followers. In an initial analysis of 14 of these candidates (excluding John Kasich, who had not yet declared his candidacy), only activity on Twitter from on or before 04 July 2015 was considered. For this period, the users who followed each candidate’s Twitter account were collected. Of these users, those who followed fewer than 13 of the candidates were discarded, leaving 1744 users. Of these remaining users, those who replied to or retweeted fewer than half of the candidates were discarded, leaving 214 users. The 50 of these users who replied to or retweeted candidates the most were selected. This selection procedure was carried out in order to identify a set of users who actively engaged in candidate-related discussion on Twitter. We chose the dimensionality of this final set of users so that a reasonably sized 50-dimensional DC pattern could be constructed, but we did not evaluate whether this dimensionality was optimal. After this initial selection procedure, a 50-dimensional vector was then calculated for each candidate based on his or her influence over each of the selected users. The influence of a candidate over a Twitter user was defined here as the number of times that user replied to or retweeted the candidate over the time period of interest.

We then performed a representational similarity analysis (RSA) as described by Kriegeskorte and colleagues (Kriegeskorte et al., 2008), in which we compared the candidates’ patterns of Twitter influence to five other measures. The first of these was a multidimensional measure of the candidates’ political positions, based on data compiled by the website ontheissues.org (retrieved 16 July 2015; Table S1). This measure recorded a candidate’s degree of support or opposition to 20 political issues based on public record (abortion, requirements to hire women and minorities, same sex marriage, public displays of God, expansion of the Affordable Care Act, privatizing Social Security, school vouchers, environmental regulations, strict punishment for crimes, rights to gun ownership, taxing the wealthy, citizenship for illegal immigrants, free trade, US sovereignty, military expansion, easing voter registration rules, foreign intervention, green energy, marijuana, and economic stimulus programs). The second measure recorded the mean polling numbers of the candidates based on five national polls conducted in late June 2015 (Table S2). The third measure was a market prediction of each candidate’s election probability from the website predictit.org (retrieved 17 July 2015; Table S3). The fourth measure was the geographical location of the candidate’s base (Table S4). The fifth measure was the candidate’s age (Table S5).

In the initial step of the RSA, we used Spearman correlation to calculate a dissimilarity matrix (DSM) for the candidates based on their patterns of Twitter influence computed above. This DSM represented the similarity between each pair of candidates based on their influence over the selected users on Twitter. We then computed DSMs for each of the five other measures. For political position, we used Spearman correlation distance. For the four remaining univariate measures, we used Euclidean distance. These DSMs represented the dissimilarity between each pair of candidates based on each of these four measures (e.g. a low value for a pair of candidates in the age DSM indicates that those two candidates were of similar age). In the final step of the RSA, the Twitter influence DSM was then correlated with each of the other measures’ DSMs. This step allowed us to evaluate whether the informational structure of candidates’ relationships based on Twitter influence matched the informational structures of their relationships based on the other measures.

A second analysis repeated the procedure above for Twitter activity in between 04 July 2015 and 12 October 2015. This analysis included John Kasich, who by then had declared his candidacy, and excluded Rick Perry, who had dropped out of the race. We also used updated measures of polling percentages (Table S2), market predictions (Table S3), and age.

## Results

### fMRI Simulation

The simulated experiment follows a standard block design, in which 15 subjects each complete 10 scanning runs consisting of five repetitions each of two conditions (“A” and “B”) interleaved with rest periods. The goal of such experiments is to identify brain activity that differs systematically between the two conditions. Results are summarized in Figure Figure 2A describes how our simulated data are constructed using a design with parameter values that are representative of real-world fMRI experiments. Figure 2C presents the results of this simulation. The classifier was able to use DC patterns to predict the condition well above the chance level of 50% [mean 67.7% accuracy (SEM 0.0223); *t*(14) = 7.93, *p* = 7.63 x 10–^7^], even with a relatively low SNR of 0.3. In contrast, none of four standard methods currently used to analyze neuroimaging data showed any difference between the two conditions. Because the signals do not vary in amplitude across conditions, a univariate analysis in which the mean signal is compared between conditions across all subjects showed no significant effect for either ROI [for source: *t*(14) = 0.575, *p* = 0.574; for destination: *t*(14) = -1.13, *p* = 0.276]. Likewise, because no multivariate activity pattern difference exists between conditions in either ROI individually, standard ROI-based multivariate classification analyses failed to distinguish between the conditions [for source: mean 54.3% accuracy (SEM 0.0345); *t*(14) = 1.13, *p* = 0.115; for destination: mean 52.0% accuracy (SEM 0.0337); *t*(14) = 0.593, *p* = 0.281]. Finally, because the overall magnitude of the influence of the source space over the destination space is constant during the experiment, existing directed connectivity measures such as MVGC and MVTE also failed to distinguish between the conditions [for MVGC: *t*(14) = 0.0389, *p* = 0.970; for MVTE: *t*(14) = -0.746, *p* = 0.468].

Figures 3A and 3B show the results of an analysis comparing the sensitivity of MDCA to the other four methods at a range of SNRs, with 100 simulations run at each SNR value. For SNR values as low as 0.2, MDCA reliably detects differences between the two conditions, while no other method detects these differences at any SNR value. Note additionally that the false positive rate for MDCA (i.e. at SNR = 0) is at the 5% level expected for our significance threshold choice of p = 0.05. Thus, MDCA provides a highly sensitive analytical tool to reveal network interactions which occur independently of phenomena such as within-region processing or average connectivity levels that are targeted by existing techniques.

To further investigate the sensitivity of MDCA, we conducted additional simulations of the above experiment to determine the minimum number of subjects, number of runs, number of data samples per DC matrix estimate, and causal strength between source and destination needed at various SNRs in order to achieve significant results. Results of these simulations are presented in Figure 3C–F. Each plot shows, for a range of SNR values, the minimum parameter value needed to achieve significance (expected *p* ≤ 0.05), while holding all other parameters fixed at the values from our first simulation. These simulations reveal that, even at SNRs below 0.3, significant results can be achieved using reasonable parameter values that could easily be incorporated into real-world experimental designs.

Figure 3G shows the results of a final simulation in which we evaluated classification accuracy in the above experimental design for a range of block durations (30 simulations per value), holding the total amount of data collected per DC matrix estimate fixed at 60 samples. Analysis results were stable whether conditions were presented in long, contiguous blocks, as is traditionally advised for directed connectivity analyses, or as isolated samples, as occurs in event-related experimental designs. A linear regression analysis of these simulation results showed no significant relationship between block duration and classification accuracy [*F*(1,358) = 0.475, *p* = 0.491]. Thus, our simulations suggest that MDCA is applicable to a range of existing experimental designs. In particular, resolving these network interactions does not appear to depend on whether data are collected contiguously or spread over many small, noncontiguous blocks.

### fMRI dataset

Our first real-world application of MDCA comes from an fMRI experiment that implemented a straightforward extension of the standard block design used in the above simulation. Figures 4A and 4B present a schematic of the experiment, which replicates an earlier investigation into visual imagery manipulation (Schlegel et al., 2016). In a series of trials over 15 scanning runs, subjects imagined one of four abstract visual shapes and performed one of four mental operations on that shape (Figure 4A).

Each trial was labeled based on either the imagined shape or the mental operation that was performed, and MDCA was performed between each pair of ROIs for each labeling scheme. Figure 4D presents the analysis results. Each arrow represents a significant DC classification result, false discovery rate (FDR) corrected for multiple comparisons across the 30 directed connections for each analysis. Dashed arrows were significant but did not pass FDR correction. The thickness of each arrow represents the effect size of the *t*-test. The analyses reveal distributed but distinct patterns of network interactions that support information about both mental representations (red arrows) and mental manipulations (blue arrows). In particular, information about mental representations was supported by bidirectional communication pathways between every pair of ROIs that we tested.

To evaluate the reliability of our MDCA method, we retested all 40 participants one month after their initial session. Figure S1 shows the results of the same analysis applied to this second dataset. The vast majority of connections appearing in Figure 4D remained significant in this new analysis. Across all 60 tests (30 directed ROI pairs and two classification labeling schemes), we found a highly significant correlation between the first and second session results (Figure 4E; [*r* = 0.769, *t*(58) = 9.16, *p* = 2.55×10^-13^]). This replication further validates our MDCA technique, showing high test-retest reliability for these data.

Because of the low temporal resolution of fMRI data, our findings were limited to interactions that occurred on the scale of seconds. MDCA applied to data with millisecond temporal resolution might have revealed further communication pathways to which fMRI data are insensitive. To evaluate this possibility, we next applied MDCA to an EEG dataset.

### EEG dataset

Our second real-world application comes from a replication of an EEG study by Donchin and colleagues of the neural basis of action preparation (Donchin et al., 1972). In our replication of this experiment, participants completed a series of trials in which they were cued to an upcoming imperative to perform an action, waited for a period of time until the imperative stimulus appeared, and then performed the action (see Figure 5A for a trial schematic).

Our question here was whether the posterior-anterior (forward) and/or the anterior-posterior (backward) directed connectivity profile of the cortex depended on the particular action that the participant was preparing for or performing. Thus, our two “nodes” in this analysis were a group of the 10 most posterior EEG electrodes and a group of the 10 most anterior electrodes (Figure 5B). Each electrode group served as both the source and the destination space, depending on the direction of connectivity being investigated, and our DC matrix estimates were constructed based on the Granger-causality from each source component to each destination component, after data were PCA transformed independently for each electrode group. Because of the high temporal resolution of EEG data, we were able to estimate these DC matrices at several time windows throughout each trial.

Similarly to the previous datasets, we asked whether we could use patterns of directed connectivity to classify which of the four actions were being prepared or performed. Figure 5C presents the results of both forward and backward directed connectivity analyses. Each two-dimensional plot shows classification accuracy as a function of window start time and lag value. As can be seen in the warm colored regions of this plot, the classifier showed highest classification accuracy for windows that included data at or after the appearance of the imperative stimulus (t = 0), indicating that action information peaked in both forward and backward cortical connectivity profiles as that action was performed. However, the classifier performed significantly above chance in every test, including windows that started at the beginning of the warning period three seconds before the imperative stimulus to perform an action (chance level was 25%, while the lowest classification accuracies were around 35%). These results indicate that the cortical connectivity profiles in each direction reflected the action even during the preparation period, when participants were performing no overt action. Finally, classification results were robust regardless of the lag value used, suggesting that the action-specific influence by each electrode group over the other extended through a range of time.

To compare how our multivariate directed connectivity method performed relative to existing directed connectivity measures, we next conducted a more traditional analysis in which we asked whether mean forward or backward connectivity depended on the action performed. Since we saw little effect of lag in our previous analysis, here we used only a single lag value of 50ms (we did not evaluate other values). Figure 5D shows the time course of classification accuracy for this lag value, taken from the analysis in Figure 5C. Figure 5E shows mean time courses of MVGC and MVTE for each of the four action types individually. At no point in these time courses did a one-way ANOVA find a significant difference between the four conditions. Figure 5F presents *p*-value time courses for each analysis, showing that the multivariate directed connectivity analysis (blue) is above the significance threshold (light blue line) at all points tested, while neither of the two traditional directed connectivity measures (red and orange) reach the threshold at any point. In other words, as far as traditional directed connectivity methods can resolve, neither forward nor backward connectivity in the brain were modulated by action type. Interestingly, MVGC does not even show that connectivity is modulated when the action is performed (see the absence in Figure 5E of a peak in the MVGC time courses around t = 0).

However, MDCA finds that patterns of both forward and backward connectivity in the cortex are modulated by the specific action that is prepared and performed. The high sensitivity of the novel method used here suggests that it may prove useful in the development of future brain-computer interfaces.

### Twitter dataset

The previous two datasets showed that MDCA can be used to investigate complex, directed patterns of connectivity in neural networks. However, nothing about these methods necessarily restrict their application to questions about the brain. To demonstrate the generality of MDCA, our final dataset comes from a quite different domain and uses an entirely different definition of directed connectivity. As of October 2015, each of the 14 Republican presidential candidates in the 2016 primary season maintained an active Twitter presence. Here we ask whether their patterns of influence over other Twitter users predict any of several variables relevant to the election cycle.

From 4 July 2015 to 12 October 2015, there were 4,231 users who followed at least 13 of the 14 declared Republican candidates on Twitter, and these followers wrote messages or “tweeted” 1,454,314 times using the service. 19,368 of these messages were replies to or “retweets” of tweets by Republican candidates (example shown in Figure 6A). Here we define our measure of directed connectivity from Twitter user *A* to user *B* to be the number of times *B* replied to or retweeted *A* over a given period of time. In general, we can compute this measure from user *A* to a range of other users in order to construct the pattern of influence of user *A* over a set of other Twitter users (Figure 6B). We did this for each of the 14 candidates, first selecting from the above set of followers only those users who had replied to or retweeted at least half of the candidates, and then taking as our destination space the 50 of these users who made the highest total volume of replies to or retweets of the candidates. Thus, for each candidate we constructed a 50-dimensional pattern of their influence over highly active Twitter users.

**Fig. 6.**
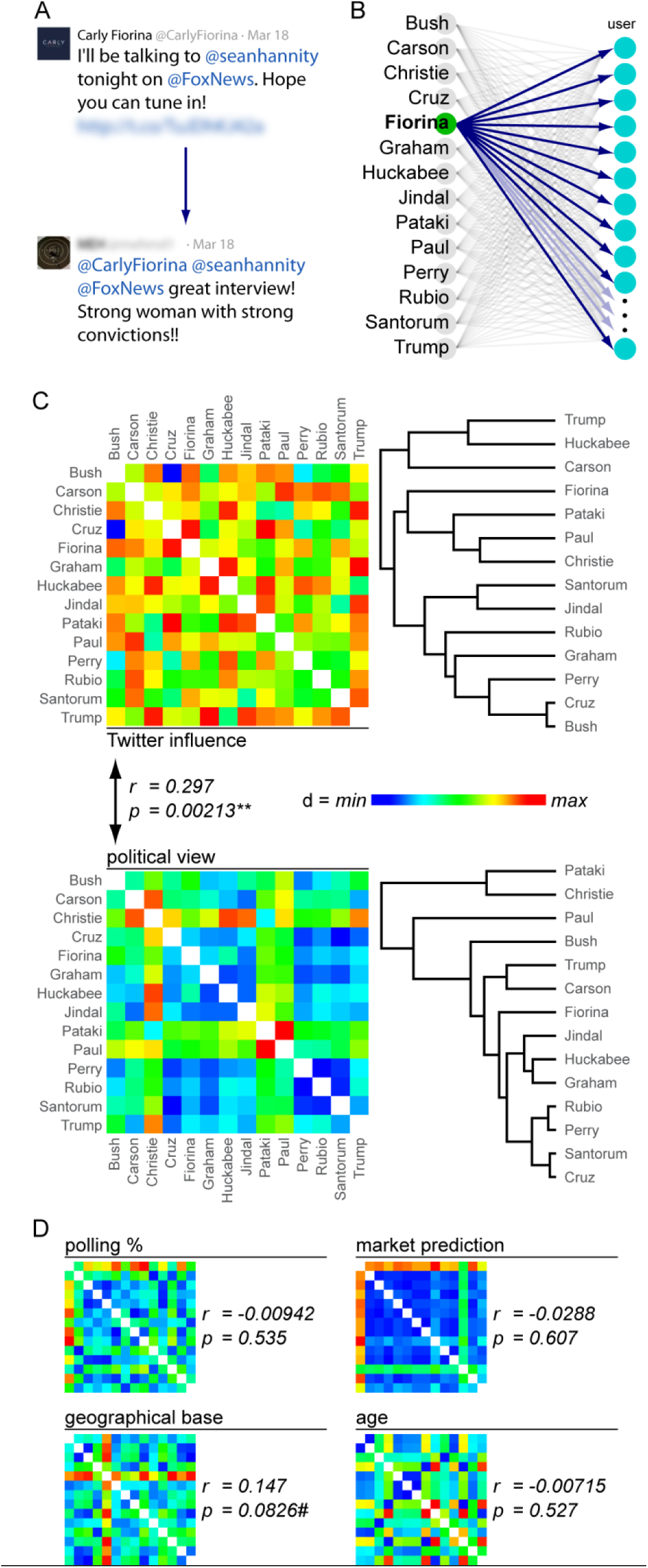
Data from a real-world study of the 2016 Republican U.S. presidential candidates’ influence on Twitter. **A.** Example of a tweet by candidate Carly Fiorina, followed by a reply from one of her followers. Directed connectivity here is defined as the number of replies by a follower over a given time period. **B.** Data for the analysis here were DC patterns from each candidate to a group of 50 shared Twitter followers. **C.** Analysis of relationship between Twitter influence and political view. At top left is a dissimilarity matrix (DSM) based on the Spearman correlation distance between the candidates’ patterns of Twitter influence. Right of that is a dendrogram representing a hierarchical clustering of candidates based on this influence. Below is a DSM and dendrogram based on a multidimensional measure of political view. Coloring is scaled independently for each DSM. These two DSMs were significantly correlated. **D.** DSMs for four other measures (candidate’s polling percentage, market prediction of candidate’s election probability, geographical location of candidate’s base, and candidate’s age), along with results of correlation analyses between these DSMs and the Twitter influence DSM.

We performed our initial analysis using only tweets made before 4 July 2015 and selecting our space of followers by applying the procedure described in Materials & Methods to these tweets. This period represented a relatively quiet time in the election cycle, before many people had begun to pay attention to the presidential election or the field of candidates. We therefore hypothesized that the character of interactions on Twitter during this period would differ qualitatively from later periods in which the campaign experienced greater public attention and media coverage. Following a standard representational similarity analysis procedure, we first used Spearman correlation distance to compute a dissimilarity matrix (DSM) between the candidates based on their estimated patterns of Twitter influence constructed as described above. This DSM is presented at the top of Figure 6C and represents the collective relationships among the candidates based on their influence on Twitter. Cool colors reflect small distances (i.e. highly similar patterns of influence) and warm colors reflect large distances (i.e. highly dissimilar patterns of influence). The dendrogram to the right shows a hierarchical clustering of the candidates based on these Twitter influence-based relationships.

Next, we constructed DSMs in a similar manner for five other measures. The first was a multidimensional measure of each candidate’s political views based on data compiled by the website ontheissues.org (retrieved 16 July 2015; Table S1). This measure recorded a candidate’s degree of support or opposition to 20 political issues (see Materials & Methods). The second measure recorded the mean polling numbers of the candidates based on five national polls conducted in late June 2015 (Table S2). The third measure was a market prediction of each candidate’s election probability from the website predictit.org (retrieved 17 July 2015; Table S3). The fourth measure was the geographical base of the candidate (Table S4). The fifth measure was the candidate’s age (Table S5). DSMs for each measure are shown in Figure 6C and 6D.

To evaluate whether any of these five measures predicted a candidate’s Twitter influence, we tested for correlations between the Twitter influence DSM and the DSMs derived from the measures (lower diagonal of each matrix only). The Twitter influence DSM correlated significantly with the political view DSM [*r* = 0.297, *t*(89) = 2.93, *p* = 0.00213], indicating that the candidates’ patterns of influence on Twitter related significantly to their political positions. The DSM based on each candidate’s geographical base also showed a marginally significant positive correlation with Twitter influence [*r* = 0.147, *t*(89) = 1.40, *p* = 0.0826]. No other correlations were significant.

However, when we performed this analysis again using data from the later period of 4 July 2015 to 12 October 2015, we no longer found a significant correlation between Twitter influence and political position [*r* = -0.0461, *t*(89) = -0.436, *p* = 0.668]. However, we did now observe a significant correlation between Twitter influence and polling numbers [*r* = 0.186, *t*(89) = 1.79, *p* = 0.0385]. Market predictions now also showed a marginally significant positive correlation with Twitter influence [*r* = 0.138, *t*(89) = 1.31, *p* = 0.0968]. These results suggest that candidate-related discourse on Twitter may have become less influenced by discussion of policy issues and more influenced by electability of the candidates as the election cycle became more active. Note that one candidate (John Kasich) entered the race and one candidate (Rick Perry) dropped out of the race in between the two analyses. All analysis results are presented in Table S6.

## Discussion

Here we have introduced a new class of methods, termed multivariate directed connectivity analyses (MDCA). We validated the method using a simulated fMRI dataset and applied it successfully to three real-world datasets (fMRI, EEG, and Twitter), showing MDCA to yield highly sensitive and repeatable effects that no previously existing analytical method can provide. We additionally showed that MDCA can be used to help resolve scientific debates concerning the role of network interactions in cognition, may be useful in the development of technologies such as brain-computer interfaces, and applies to a range of data modalities in and beyond neuroscience.

Our simulated fMRI dataset adopted a standard block design that contrasted two experimental conditions. This type of design is used widely in the field to investigate a range of questions. Importantly, our two conditions differed only in the patterns of directed connectivity that they induced between two regions. Such situations might be expected to occur in real-world scenarios during complex cognitive processes that entail relatively constant overall mean levels of neural activity and bidirectional interregional interactions, but in which complex patterns of interaction change dynamically as a process unfolds. Thus, while existing methods such as univariate directed connectivity and ROI-based multivariate classification analyses could not detect any differences between the neural activity induced by the two conditions, MDCA was highly sensitive to differences between the two conditions based on changes in the patterns of directed connectivity that occurred. In further simulations, we showed that MDCA is highly sensitive even at low SNRs and works for a range of designs and parameter values used in typical fMRI experiments.

Note that our simulation was simplified in several important respects from real-world fMRI data. We did not: convolve our generated data with an HRF; correct for that HRF when analyzing the simulated data; construct our data in a spatially realistic manner; simulate movement artifacts; or simulate noise processes based on empirical measurements of fMRI data. This simulation was not intended to provide a high-fidelity evaluation of MDCA’s applicability to fMRI data specifically, but instead was intended to evaluate in a general manner whether the method could recover multivariate patterns of directed connectivity from time series data when other methods failed. This choice to construct a simple but general evaluation of the method using simulated data makes its evaluation on real-world fMRI (and other) data necessary.

The analyses of our real-world fMRI data were able to probe the distributed nature of information processing in a cortex-wide network to a degree of specificity not previously possible. Specifically, we showed that both the representation and manipulation of mental visual images entails a dense network of bidirectional connectivity throughout the cerebral cortex. Several standard models of working memory postulate that it is mediated by a network of informationally segregated anatomical modules, including a “central executive” that resides in DLPFC and a “visuospatial sketchpad” that resides in posterior visual cortex (Baddeley, 2003; Postle, 2006). However, MDCA reveals evidence that information about both “executive” and “sketchpad” aspects of a working memory task occurs in a dense flow of information between many overlapping cortical regions. This finding calls for more distributed models of working memory and is consistent with recent work in monkeys showing that information quickly becomes highly distributed in the brain during cognitive tasks (Siegel et al., 2015).

Of particular note is the high correlation between analysis results in our initial and followup scans of each of the original 40 participants. This high test-retest reliability further validates MDCA as a technique that can be used in practice to analyze fMRI data (Figure S1).

Note, however, that fMRI places constraints on the processes that can be studied using functional connectivity-based analyses. Because of its low temporal resolution, fMRI is limited to resolving processes that play out over a number of seconds, as in the visual imagery manipulation task used here. In other words, MDCA applied to fMRI data cannot be used to resolve information transfer at the level of neuron-to-neuron communication. Indeed, no technique could accomplish this even in theory when applied to fMRI data, since the blood flow-related changes measured by fMRI are fundamentally too coarse for such resolution. Rather, with fMRI data, MDCA should be limited to the investigation of distributed processes that occur far higher in the hierarchy of information processing and that play out over seconds rather than milliseconds. The key point here is that directed causal interactions in the cortex occur at a range of timescales, from milliseconds for low-level neuronal interactions, to potentially seconds (or years) at higher levels of the information processing hierarchy, and different techniques will be sensitive to different scales of interaction. Thus, researchers planning to use MDCA in their research should consider the temporal characteristics of the phenomena they wish to study.

EEG data, however, do not suffer from the temporal limitations of fMRI. To illustrate, our EEG analysis revealed that information about the preparation of specific actions is accessible to MDCA in a time-resolved way, potentially starting seconds before an action is performed. Existing, univariate measures of directed connectivity were insensitive to these action-preceding processes. Our analysis suggests that MDCA can be combined with existing technologies to provide a powerful tool to predict and respond to cognitive processes that precede action. This technique could be used, for instance, to predict the intentions of patients and use these predictions to control a range of instruments including computer mouse cursors or artificial limbs (Nicolelis, 2001).

We used Granger-causality (Barnett and Seth, 2014) as a directed connectivity measure in our fMRI and EEG datasets. It should be noted, however, that Granger-causality does not measure causation directly. Instead, Granger-causality is a statistical method for evaluating the ability of a source signal to predict the future of a destination signal beyond the predictive power provided by the destination signal’s own past. While the validity of Granger-causality for fMRI data recently came under scrutiny, subsequent computational and empirical work has shown that it is a viable technique when proper precautions such as those used in the present study are taken (Barnett and Seth, 2014; Friston et al., 2013; Wen et al., 2013). In particular, we investigated differences in patterns of Granger-causality between conditions rather than attempting to establish “ground-truth” connectivity between regions. However, when MDCA employs directed connectivity measures based on statistical predictability rather than causality, it cannot be used to make strong conclusions about causation per se. In situations in which direct measures of causality are available (such as our Twitter dataset), MDCA can then make stronger causal conclusions by using those causal measures rather than statistical measures such as Granger-causality or transfer entropy.

Finally, our Twitter dataset demonstrated that MDCA can reveal the underlying informational structure of complex patterns of interactions in a range of networks. Our data suggest that the 2016 Republican presidential candidates’ influence on the social media network Twitter was shaped initially by their political positions and later by their electability. Our analysis of these data additionally demonstrated that MDCA can be applied using a wide range of definitions of directed connectivity and a large class of existing multivariate methods. Thus MDCA may be applicable to a range of fields in biology, computer science, Earth science, and beyond that demand the characterization of complex network interactions.

While many important phenomena in nature arise out of complex interactions within information networks, the empirical study of such network dynamics has until recently suffered from both a lack of tools to collect data about these networks and a lack of analytical methods to study what data were available. The recently launched BRAIN Initiative in the U.S. attempts to address this issue, largely through the development of new tools for measuring brain activity and connectivity at unprecedented levels of spatial and temporal detail (Insel et al., 2013). However, the increase in our capacity to measure the brain’s complexity brings with it a new class of “Big Data” (Turk-Browne, 2013) problems for neuroscience: How do we make sense of all these data? In other words, how can we translate measurements of the structure and dynamics of a complex network such as the brain into a mechanistic understanding of that network’s functions? We believe that MDCA will provide an important addition to the growing set of tools available to answer these questions.

Here we show that using MDCA to treat network interactions as multidimensional and information-rich leads to the possibility of new mechanistic insights into network dynamics. In our real-world data, MDCA provides evidence for highly distributed processing during the mental manipulation of visual imagery, can predict with high sensitivity the particular action being prepared by a subject even seconds before that action is performed, and reveals patterns of influence on the social media network Twitter that reflect an evolving presidential political landscape. Thus, multivariate directed connectivity analyses can reveal behavior in networks that traditional methods fail to detect. These methods are sensitive and reliable, can be incorporated into existing experimental designs, and apply to networks across a range of fields of study. New analytical methods such as these may prove vital to efforts such as the BRAIN Initiative that seek to understand deep, complex contemporary scientific problems including the biological basis of the mind.

## Acknowledgments

Data and code reported in the paper are available in Supplementary Materials and at http://www.dartmouth.edu/~petertse. This study was funded by NSF Graduate Research Fellowship 2012095475 to AS, the Neukom Institute, NSF Grant 1632738 to PT, and Templeton Foundation Grant 14316 to PT. AS designed the studies and developed the analytical methods. AS and BV developed the simulations and analyzed the data. AS and PA collected the data. PT supervised the project. All authors wrote the paper.

## Supplementary Figures

**Fig. S1.**
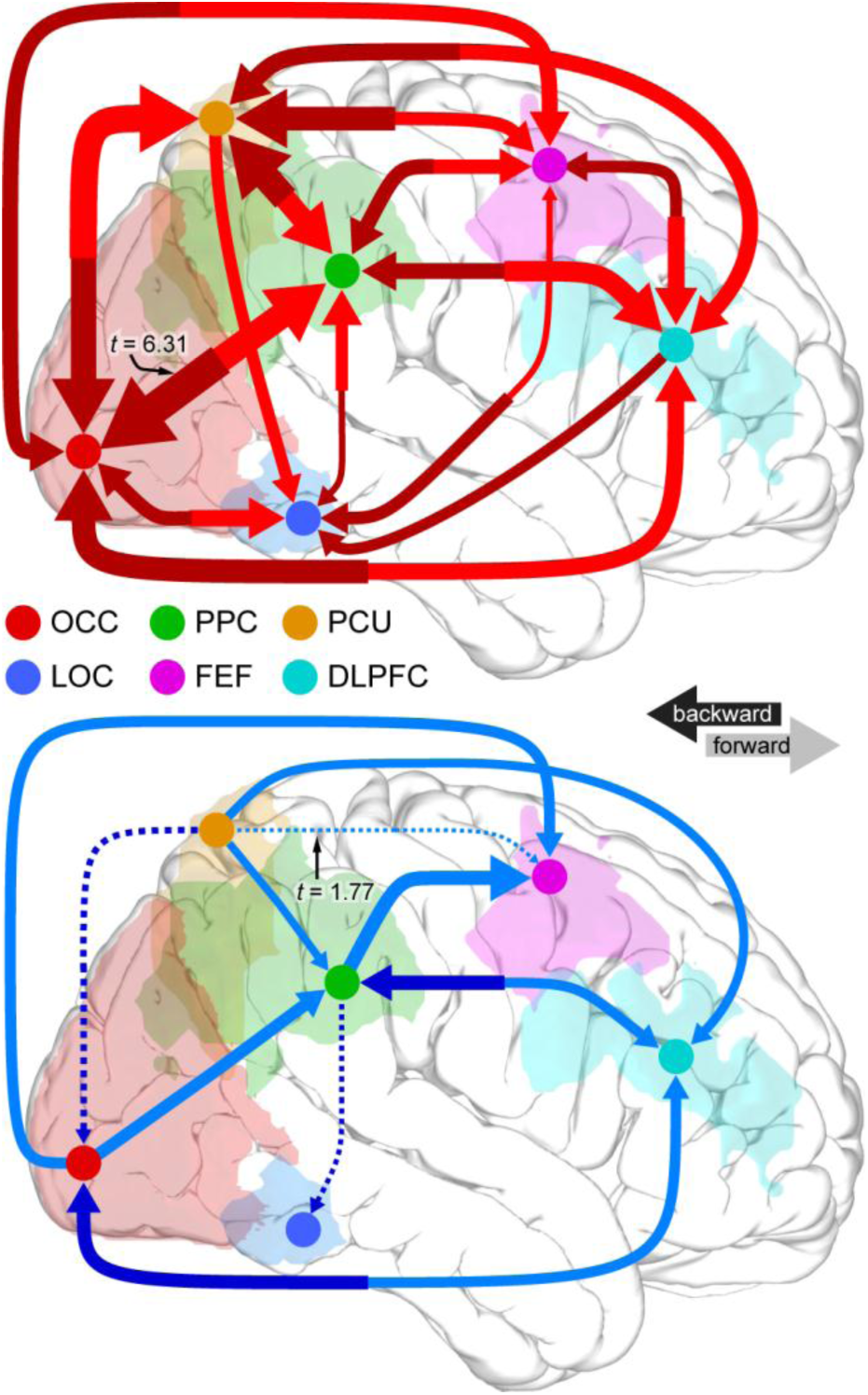
MDCA Test-retest reliability. DC classification results for one-month retest of participants from the fMRI experiment in Figure 4.

## Supplementary Tables

**Table S1.** Candidate political views, as compiled by ontheissues.org (retrieved 16 July 2015).

|  | Bush | Carson | Christie | Cruz | Fiorina | Graham | Huckabee | Jindal | Kasich | Pataki | Paul | Perry | Rubio | Santorum | Trump |
| --- | --- | --- | --- | --- | --- | --- | --- | --- | --- | --- | --- | --- | --- | --- | --- |
| abortion rights | -5 | -5 | -3 | -5 | -3 | -5 | -5 | -5 | -5 | -3 | -5 | -5 | -5 | -5 | -3 |
| require hiring of woman & minorities | 2 | 5 | -3 | 0 | -3 | 2 | 2 | 5 | -3 | -3 | 2 | 0 | 2 | -3 | 2 |
| same sex marriage | -3 | -3 | 0 | -5 | -3 | -5 | -5 | -5 | -5 | 5 | -3 | -5 | -5 | -5 | -3 |
| public displays of God | -5 | -5 | 2 | -5 | 0 | -5 | -5 | -3 | -5 | -3 | -5 | -5 | -5 | -5 | -5 |
| expand the Affordable Care Act | 2 | 2 | -3 | 5 | 5 | 2 | 2 | 5 | 2 | -3 | 5 | 5 | 2 | 5 | 2 |
| privatize Social Security | 0 | 0 | 2 | 5 | 0 | 5 | 2 | 0 | 5 | 2 | 2 | 2 | -3 | 5 | 2 |
| school vouchers | 5 | 5 | 5 | 2 | 2 | 5 | 2 | 5 | 2 | 5 | 5 | 5 | 5 | 5 | 5 |
| environmental regulation | 5 | 2 | 0 | -3 | -3 | -3 | 2 | 0 | 0 | 2 | -5 | -3 | -5 | -5 | -3 |
| strict punishment for crimes | -5 | -3 | -3 | -5 | -5 | -5 | -3 | -5 | -5 | -5 | 5 | -5 | -5 | -5 | -5 |
| rights to gun ownership | 2 | 2 | -3 | 5 | 5 | 5 | 5 | 5 | 5 | -3 | 5 | 5 | 5 | 2 | 2 |
| taxing the wealthy | 5 | 5 | 5 | 2 | 5 | 5 | 2 | 5 | 5 | 2 | 5 | 5 | 5 | 5 | 0 |
| citizenship for illegal immigrants | 2 | -5 | 2 | -5 | -3 | 2 | -3 | -5 | 0 | -3 | 2 | -3 | -3 | -5 | -5 |
| free trade | 2 | -5 | 2 | 5 | -3 | -3 | -3 | -5 | 5 | 2 | 5 | 2 | 5 | 5 | -3 |
| US sovereignty | 0 | 5 | -3 | 5 | 2 | 5 | 5 | 5 | -5 | 0 | -3 | -3 | 2 | 5 | 5 |
| military expansion | -5 | 2 | -5 | -5 | -3 | -5 | -3 | -3 | 2 | -3 | 2 | -5 | -5 | -5 | -5 |
| ease voter registration | -5 | 0 | 2 | -3 | 2 | 0 | -5 | 2 | 2 | 0 | 5 | -3 | 2 | -2 | 0 |
| foreign intervention | -3 | 2 | -3 | -3 | -3 | -5 | -5 | -5 | -3 | -3 | 2 | -5 | -5 | -5 | 2 |
| green energy | 2 | 2 | -5 | 5 | 2 | 5 | 2 | 5 | 2 | -3 | 5 | 2 | 2 | 5 | 5 |
| marijuana | -5 | -3 | 2 | 0 | 0 | -5 | -3 | -3 | -5 | -3 | 5 | -3 | -5 | -5 | 0 |
| economic stimulus | 2 | 5 | 5 | 5 | 5 | 2 | 2 | 2 | 0 | 5 | 2 | 5 | 5 | 5 | 5 |

**Table S2.** Polling percentage source and values.

|  | Bush | Carson | Christie | Cruz | Fiorina | Graham | Huckabee | Jindal | Kasich | Pataki | Paul | Perry | Rubio | Santorum | Trump |
| --- | --- | --- | --- | --- | --- | --- | --- | --- | --- | --- | --- | --- | --- | --- | --- |
| **First analysis:** |  |  |  |  |  |  |  |  |  |  |  |  |  |  |  |
| 13-15 June 2015<br>Economist/YouGov | 14 | 9 | 3 | 4 | 6 | 2 | 6 | 3 | N/A | 0 | 11 | 2 | 10 | 1 | 11 |
| 14-18 June 2015<br>NBC/WSJ | 17 | 8 | 3 | 3 | 1 | 1 | 5 | 2 | N/A | 0 | 8 | 4 | 7 | 4 | 12 |
| 20-22 June 2015<br>Economist/YouGov | 10 | 10 | 2 | 9 | 3 | 2 | 6 | 0 | N/A | 0 | 11 | 2 | 11 | 2 | 11 |
| 26-28 June 2015<br>CNN | 22 | 11 | 4 | 4 | 2 | 1 | 9 | 0 | N/A | 0 | 7 | 5 | 14 | 0 | 1 |
| 27-29 June 2015<br>Economist/YouGov | 14 | 9 | 4 | 3 | 6 | 0 | 7 | 1 | N/A | 0 | 9 | 7 | 10 | 3 | 2 |
| **Second analysis:** |  |  |  |  |  |  |  |  |  |  |  |  |  |  |  |
| 17-19 Sept. 2015<br>CNN | 10 | 17 | 2 | 7 | 6 | 1 | 4 | 1 | 4 | 1 | 2 | N/A | 13 | 2 | 27 |
| 17-21 Sept. 2015<br>Quinnipiac | 8 | 24 | 2 | 6 | 9 | 0 | 2 | 1 | 4 | 0 | 3 | N/A | 11 | 0 | 17 |
| 18-21 Sept. 2015<br>Bloomberg | 8 | 13 | 1 | 6 | 13 | 1 | 2 | 1 | 2 | 0 | 2 | N/A | 9 | 0 | 23 |
| 20-22 Sept. 2015<br>Fox News | 4 | 16 | 1 | 6 | 8 | 1 | 2 | 0 | 1 | 0 | 2 | N/A | 8 | 0 | 25 |
| 20-24 Sept. 2015<br>NBC/WSJ | 7 | 20 | 3 | 5 | 11 | 0 | 2 | 1 | 6 | 0 | 3 | N/A | 11 | 1 | 21 |
| 22-27 Sept. 2015<br>Pew | 7 | 18 | 5 | 8 | 9 | 0 | 3 | 0 | 4 | 1 | 2 | N/A | 9 | 0 | 26 |
| 24 Sept. 2015<br>USA Today | 13 | 16 | 4 | 5 | 11 | 0 | 3 | 1 | 4 | 0 | 2 | N/A | 8 | 1 | 21 |
| 26 Sept. – 2 Oct.<br>2015, Investor's<br>Business Daily | 10 | 17 | 2 | 7 | 12 | 0 | 2 | 0 | 2 | 1 | 1 | N/A | 9 | 0 | 25 |
| 1-4 October 2015<br>Public Policy<br>Polling | 9 | 14 | 3 | 6 | 15 | 0 | 6 | 0 | 2 | 0 | 4 | N/A | 11 | 1 | 24 |

**Table S3.** Market prediction of candidate election probability, according to the website predictit.org.

|  | 1<sup>st</sup> analysis<br>(retrieved 17 July<br>2015) | 2<sup>nd</sup> analysis<br>(retrieved 7<br>October 2015) |
| --- | --- | --- |
| Bush | 47 | 31 |
| Carson | 9 | 13 |
| Christie | 8 | 7 |
| Cruz | 11 | 12 |
| Fiorina | 7 | 13 |
| Graham | 8 | 2 |
| Huckabee | 10 | 4 |
| Jindal | 10 | 4 |
| Kasich | N/A | 11 |
| Pataki | 3 | 2 |
| Paul | 12 | 7 |
| Perry | 11 | N/A |
| Rubio | 27 | 40 |
| Santorum | 6 | 4 |
| Trump | 15 | 17 |

**Table S4.**
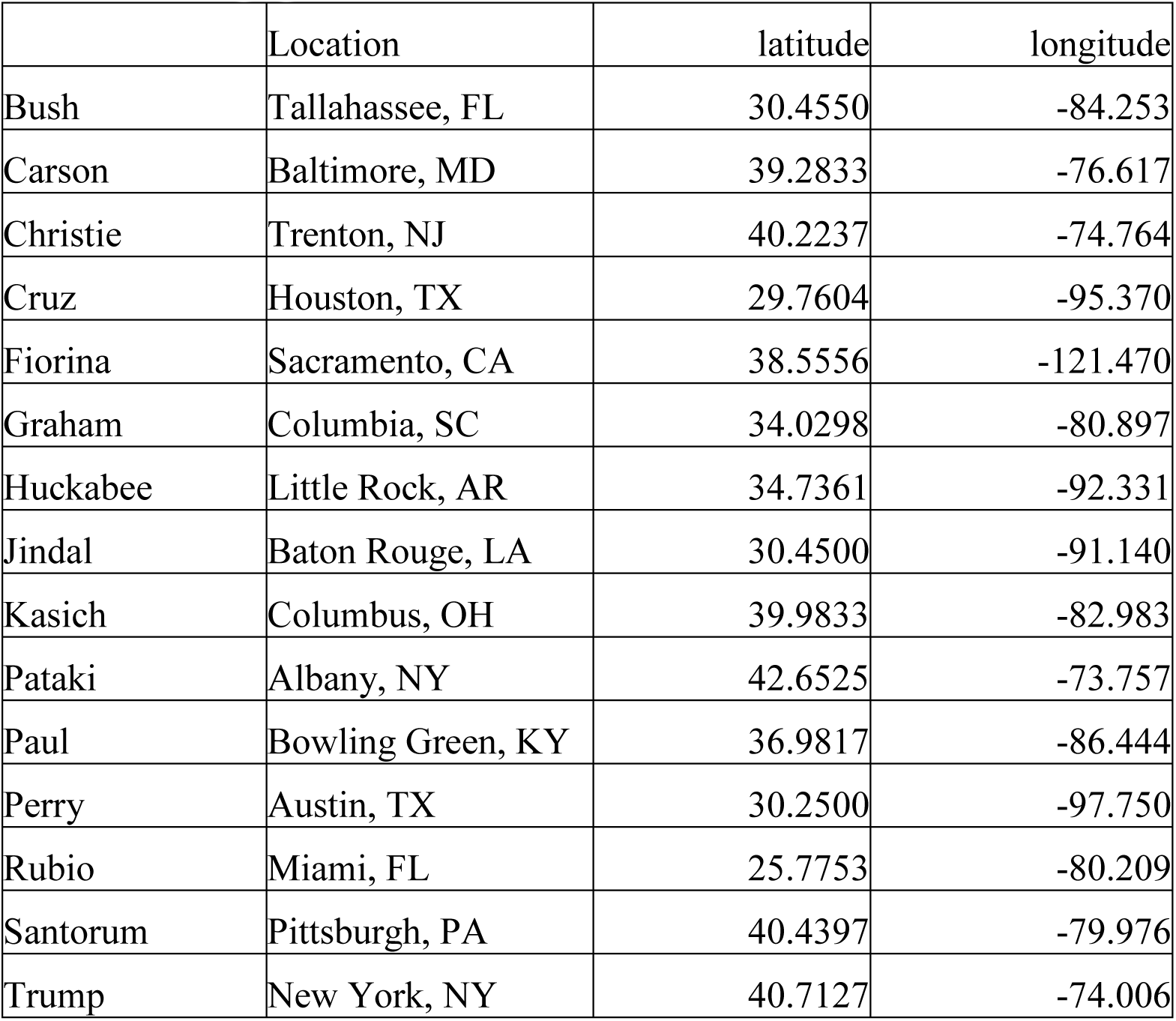
Geographical location of each candidate’s base.

**Table S5.** Candidate dates of birth.

|  | DOB |
| --- | --- |
| Bush | 11 Feb. 1952 |
| Carson | 18 Sept. 1951 |
| Christie | 6 Sept. 1962 |
| Cruz | 22 Dec. 1970 |
| Fiorina | 6 Sept. 1954 |
| Graham | 9 July 1955 |
| Huckabee | 24 Aug. 1955 |
| Jindal | 10 June 1961 |
| Kasich | 13 May 1952 |
| Pataki | 24 June 1945 |
| Paul | 7 Jan. 1963 |
| Perry | 4 Mar. 1950 |
| Rubio | 28 May 1971 |
| Santorum | 10 May 1958 |
| Trump | 14 June 1946 |

**Table S6.**
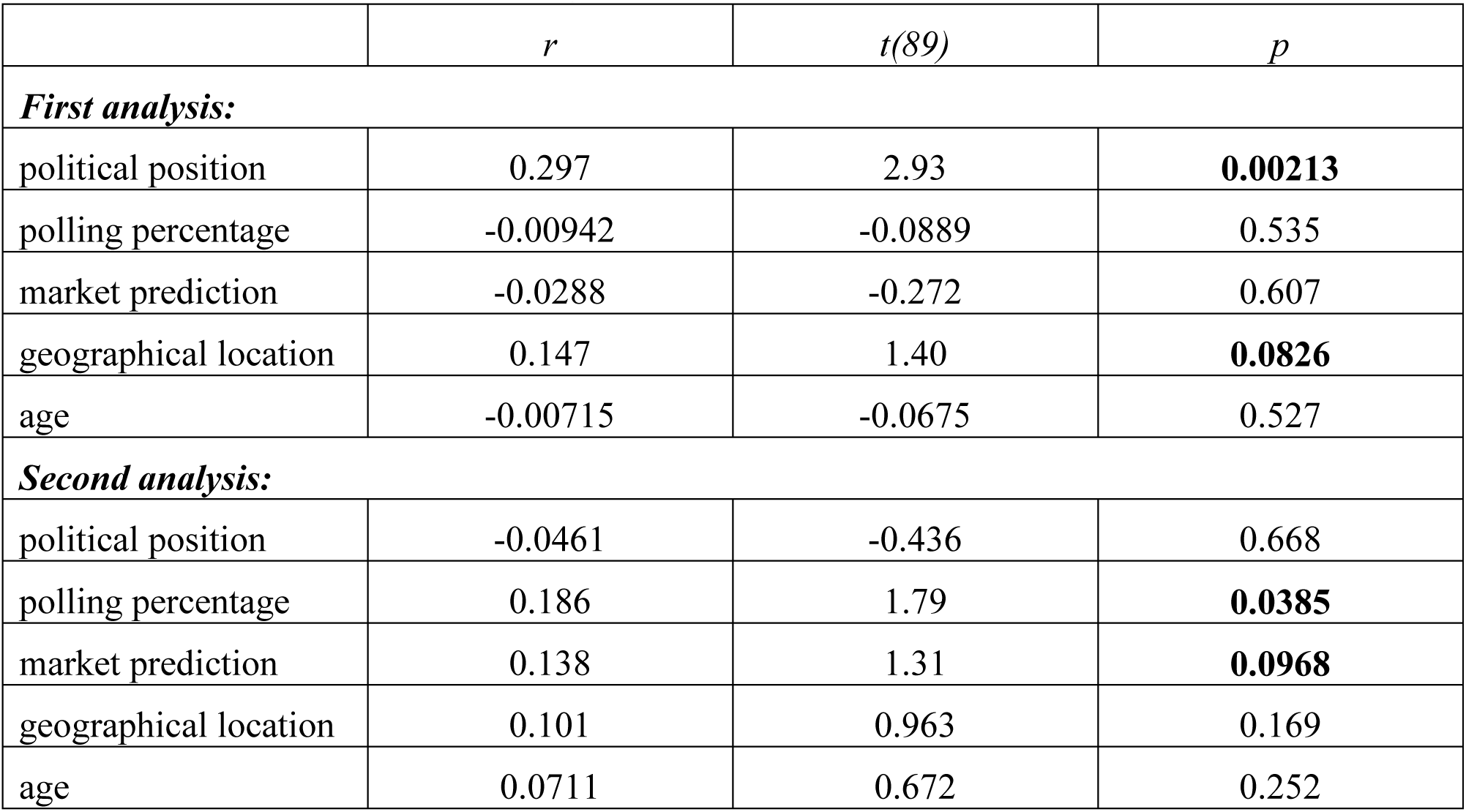
Results of correlation analyses between Twitter influence DSM and other DSMs.

|  | *r* | *t*(89) | *p* |
| --- | --- | --- | --- |
| ***First analysis:*** |  |  |  |
| political position | 0.297 | 2.93 | **0.00213** |
| polling percentage | -0.00942 | -0.0889 | 0.535 |
| market prediction | -0.0288 | -0.272 | 0.607 |
| geographical location | 0.147 | 1.40 | **0.0826** |
| age | -0.00715 | -0.0675 | 0.527 |
| ***Second analysis:*** |  |  |  |
| political position | -0.0461 | -0.436 | 0.668 |
| polling percentage | 0.186 | 1.79 | **0.0385** |
| market prediction | 0.138 | 1.31 | **0.0968** |
| geographical location | 0.101 | 0.963 | 0.169 |
| age | 0.0711 | 0.672 | 0.252 |

**Table S7.**
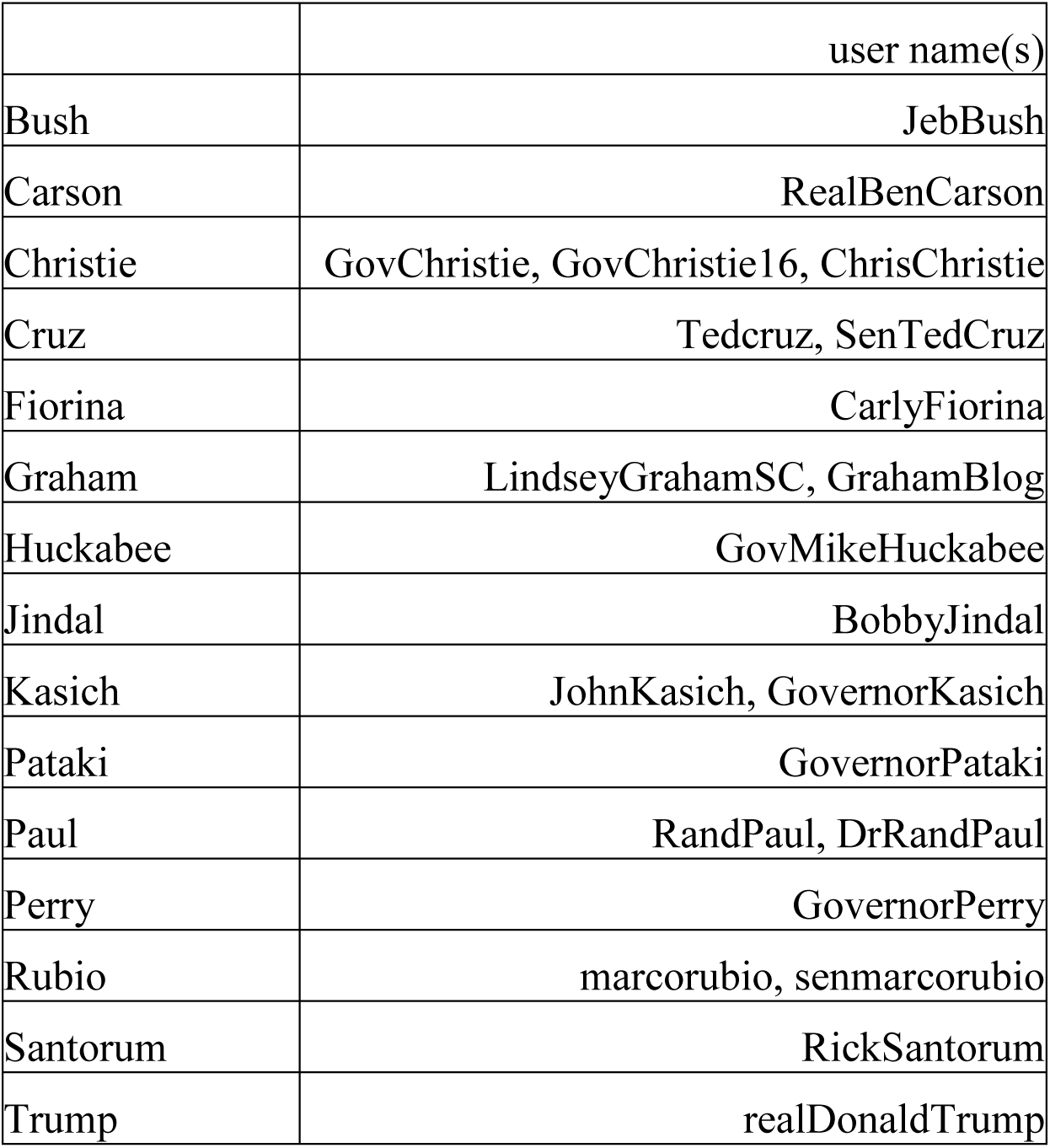
Candidates’ Twitter user names.

|  | user name(s) |
| --- | --- |
| Bush | JebBush |
| Carson | RealBenCarson |
| Christie | GovChristie, GovChristie16, ChrisChristie |
| Cruz | Tedcruz, SenTedCruz |
| Fiorina | CarlyFiorina |
| Graham | LindseyGrahamSC, GrahamBlog |
| Huckabee | GovMikeHuckabee |
| Jindal | BobbyJindal |
| Kasich | JohnKasich, GovernorKasich |
| Pataki | GovernorPataki |
| Paul | RandPaul, DrRandPaul |
| Perry | GovernorPerry |
| Rubio | marcorubio, senmarcorubio |
| Santorum | RickSantorum |
| Trump | realDonaldTrump |

